## Supplemental information for "Pinging the Hidden Attentional Priority Map: Suppression Needs Attention"

### **Supplementary Information**

**Figure S1**

*No Evidence of Behavioral Interaction Between Spatial Working Memory and Visual Search*

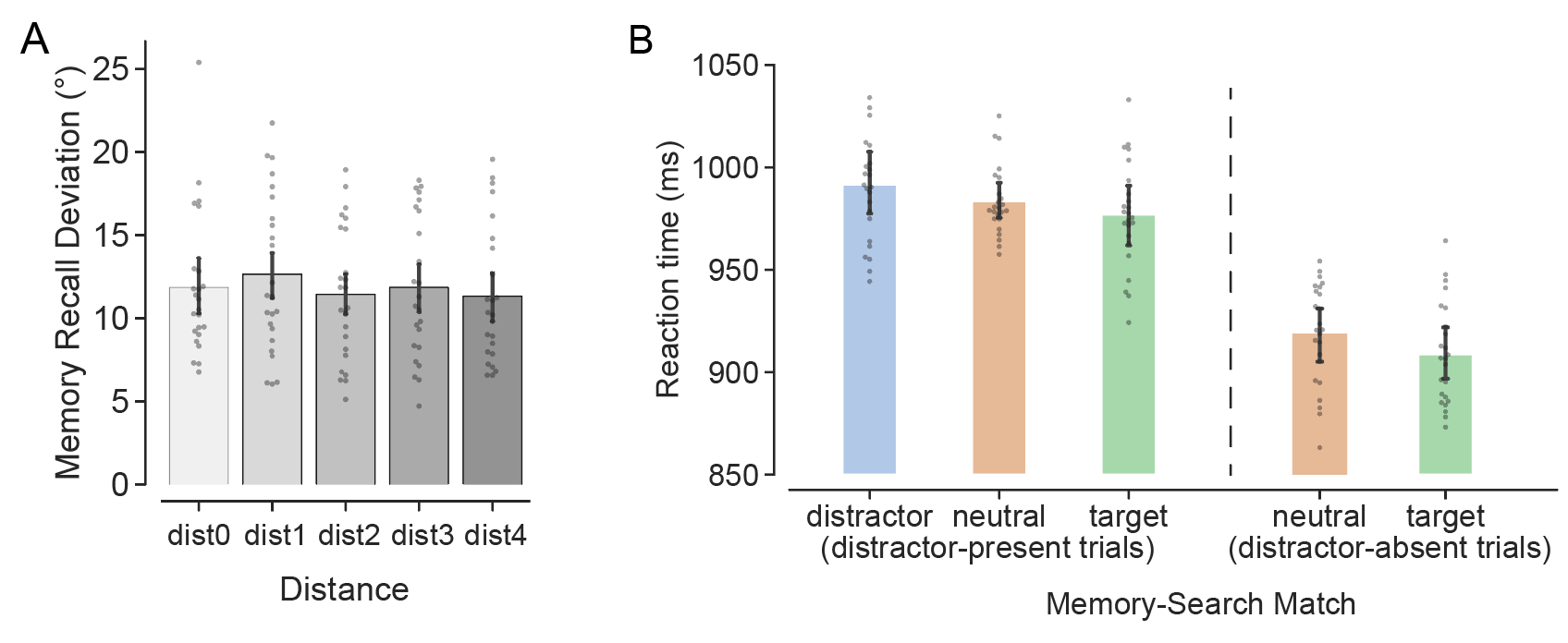

*Note.* (A) The average memory recall deviation as a function of the distance between memory cue and high-probability location. (B) The average reaction times as a function of the spatial overlap between the memory cue location and any of the search items, separating trials by distractor-present (match-target, match-distractor, match-neutral) and distractor-absent (match-target, match-neutral) conditions.

To investigate the potential interactions between the spatial memory task and the visual search task, we conducted additional analyses on the behavioral data. First, we examined whether memory recall was influenced by the spatial distance (dist0 to dist4) between the memory cue location and the high-probability distractor location. As shown in the Figure S1A, memory recall is not systematically biased either toward or away from the high-probability distractor location (p = .562, ηp² = .011).

To assessed how the memory task might affect search performance, we plotted reaction times as a function of the spatial overlap between the memory cue location and any of the search items, separating trials by distractor-present (match-target, match-distractor, match-neutral) and distractor-absent (match-target, match-neutral) conditions. Although visually the result pattern (Figure S1B) seems to suggest that search performance was facilitated when the memory cue spatially overlapped with the target and interfered with when it overlapped with the distractor, this pattern did not reach statistical significance (distractor-present: p = .249, ηp² = .002; distractor-absent: p = .335, ηp² = .002).

**Figure S2**

*The performance of the Localizer in the training phase*

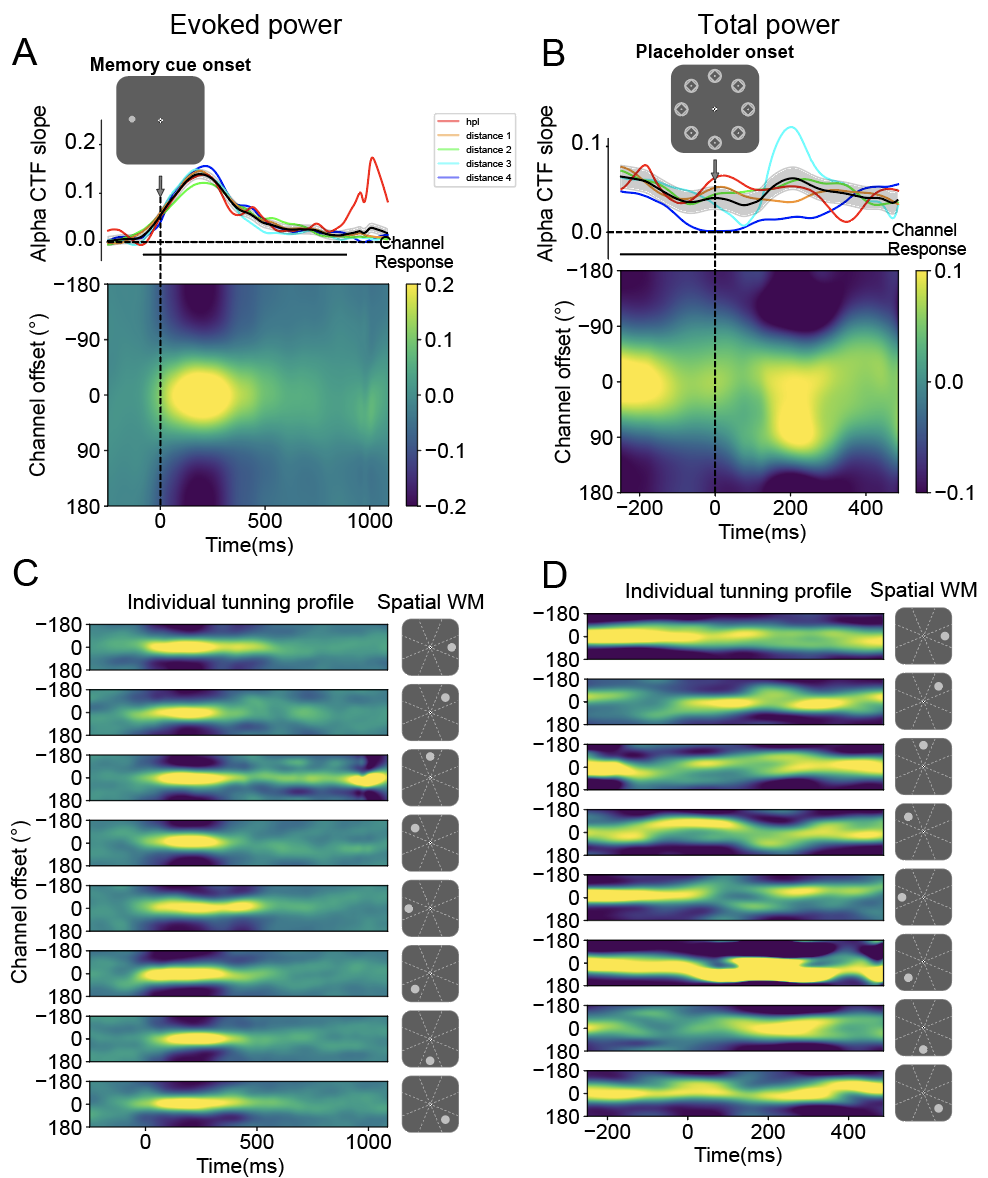

*Note.* (A) The average evoked alpha-band channel tunning function (CTF) profiles time-locked to memory onset. (B) The average total alpha CTF profiles time-locked to placeholder onset. The lower image depicts the responses across channels, while the plot above shows CTF slopes, with amplitude signifying spatial selectivity. Shaded areas reflect bootstrapped SEM. Time points exhibiting significant differences in CTF slopes, identified through a cluster-based permutation test (p < .05), are marked with horizontal black insets. The light color lines in the background indicate the CTF slopes tuned to different memory cue locations, grouped by their distance to the artificial high-probability location. (C) Individual total CTF profiles, synchronized with memory cue onset, finely tuned to each of the eight spatial cue locations, respectively. (D) Individual evoked CTF profiles, synchronized with placeholder onset, finely tuned to each of the eight spatial cue locations, respectively.
